# Memory engram synapse 3D molecular architecture visualized by cryoCLEM-guided cryoET

**DOI:** 10.1101/2025.01.09.632151

**Authors:** Charlie Lovatt, Thomas J. O’Sullivan, Clara Ortega-de San Luis, Tomás J. Ryan, René A. W. Frank

## Abstract

Memory is incorporated into the brain as physicochemical changes to engram cells. These are neuronal populations that form complex neuroanatomical circuits, are modified by experiences to store information, and allow for memory recall. At the molecular level, learning modifies synaptic communication to rewire engram circuits, a mechanism known as synaptic plasticity. However, despite its functional role on memory formation, the 3D molecular architecture of synapses within engram circuits is unknown. Here, we demonstrate the use of engram labelling technology and cryogenic correlated light and electron microscopy (cryoCLEM)-guided cryogenic electron tomography (cryoET) to visualize the in-tissue 3D molecular architecture of engram synapses of a contextual fear memory within the CA1 region of the mouse hippocampus. Engram cells exhibited structural diversity of macromolecular constituents and organelles in both pre- and postsynaptic compartments and within the synaptic cleft, including in clusters of membrane proteins, synaptic vesicle occupancy, and F-actin copy number. This ‘engram to tomogram’ approach, harnessing *in vivo* functional neuroscience and structural biology, provides a methodological framework for testing fundamental molecular plasticity mechanisms within engram circuits during memory encoding, storage and recall.

## Introduction

Memory is the process by which the brain encodes, retains, and recalls information to alter future behaviour. Engram cells, coined by Richard Semon^1^, are the hypothetical substrate of memory storage. These are operationalized as subsets of neurons that, modified upon experiences, underlie physicochemical changes to store information, and are reactivated by relevant cues to elicit memory recall and a behavioral response ^2–6^.

Within the brain, neurons are wired into complex neuroanatomical circuitry via synaptic connections that are dynamically modified by plasticity mechanisms^7–10^. Plasticity modulates synapse formation, synaptic strength, and homeostasis^8,9,11–13^ to enable learning and memory^8,9,14^. Structural plasticity mediates the enduring modification in synaptic wiring patterns, providing a plausible physical substrate for retaining long-term information^15,16^.

Engram labelling technology, based on the activity-dependent targeting of neurons associated with discrete memories, allows the identification, characterization and *in vivo* manipulation of putative engram cells^2,3,17^. The use of this set of tools demonstrated that memory encoding is associated with cellular structural changes^17–21^ via plasticity of functional connectivity of engram circuits^17,22–25^. Increases in synaptic strength of engram cell synapses correlates positively with the acquisition and retrievability of memory in learning paradigms^17–20,26^.

The structure of synapses and their change of shape after *in vitro*-evoked synaptic plasticity have been described by early pioneering conventional electron microscopy studies^27–29^. Serial sectioning using focused ion beam milling has provided a volumetric reconstruction of neurons, revealing, for example, that more synapse-dense regions are found rostrally^30^. However, these approaches are unable to resolve the native structure of individual proteins within the synapse. In contrast, cryo-electron tomography (cryoET) can capture the 3D molecular architecture of tissues in a native, vitreous state unperturbed by chemical fixation and staining methods. Various sample preparations for cryoET have been developed to investigate synapses, such as with synaptosomes^31–33^, tissue homogenate^34^, cultured neurons^35–37^ and intact tissue^34,38–40^. However, the molecular architecture of synapses within a neuronal circuit encoding a specific memory has not yet been visualized in any fresh brain tissue preparation.

Here, we developed a workflow integrating engram labelling technology^2^ with cryogenic correlated light and electron microscopy (cryoCLEM)^41^ and cryoET to determine the in-tissue 3D molecular architecture of synapses within neuronal engram circuits. We used genetically encoded fluorophores to label pre- and postsynaptic neurons of a contextual fear conditioning engram circuit, from CA3 to CA1 in the hippocampus. CryoCLEM-guided cryoET of engram synapses at this synaptic junction revealed the molecular architecture of engram synapses, including identifying and quantifying organelles and macromolecular constituents. Our study highlights the heterogeneity of synapses in engram circuits and offers a readily adaptable workflow to investigate specific molecular mechanisms and structural features within engram cells and circuits.

## Results

### CryoCLEM-guided cryoET of engram synapse

We developed a workflow to determine the in-tissue molecular architecture of engram synapses within fresh tissue using engram labelling technology, cryoCLEM and cryoET (**Fig. 1**). We focused on the *Schaffer collateral* circuit between CA3 and CA1 hippocampal neurons that allow contextual learning^42^. To label an engram encoding an aversive, context-specific fear memory, two AAVs cocktails: *c-fos-tTA*/*TRE-ChR2-EYFP* and *c-fos-tTA*/*TRE-mCherry* were intracerebrally injected into the CA3 and contralateral CA1, respectively (**Fig. 1A**). Upon removal of doxycycline from the diet, animals underwent a contextual fear conditioning paradigm (context-associated foot-shocks), forming an engram for an aversive contextual fear memory.

**Figure 1.**
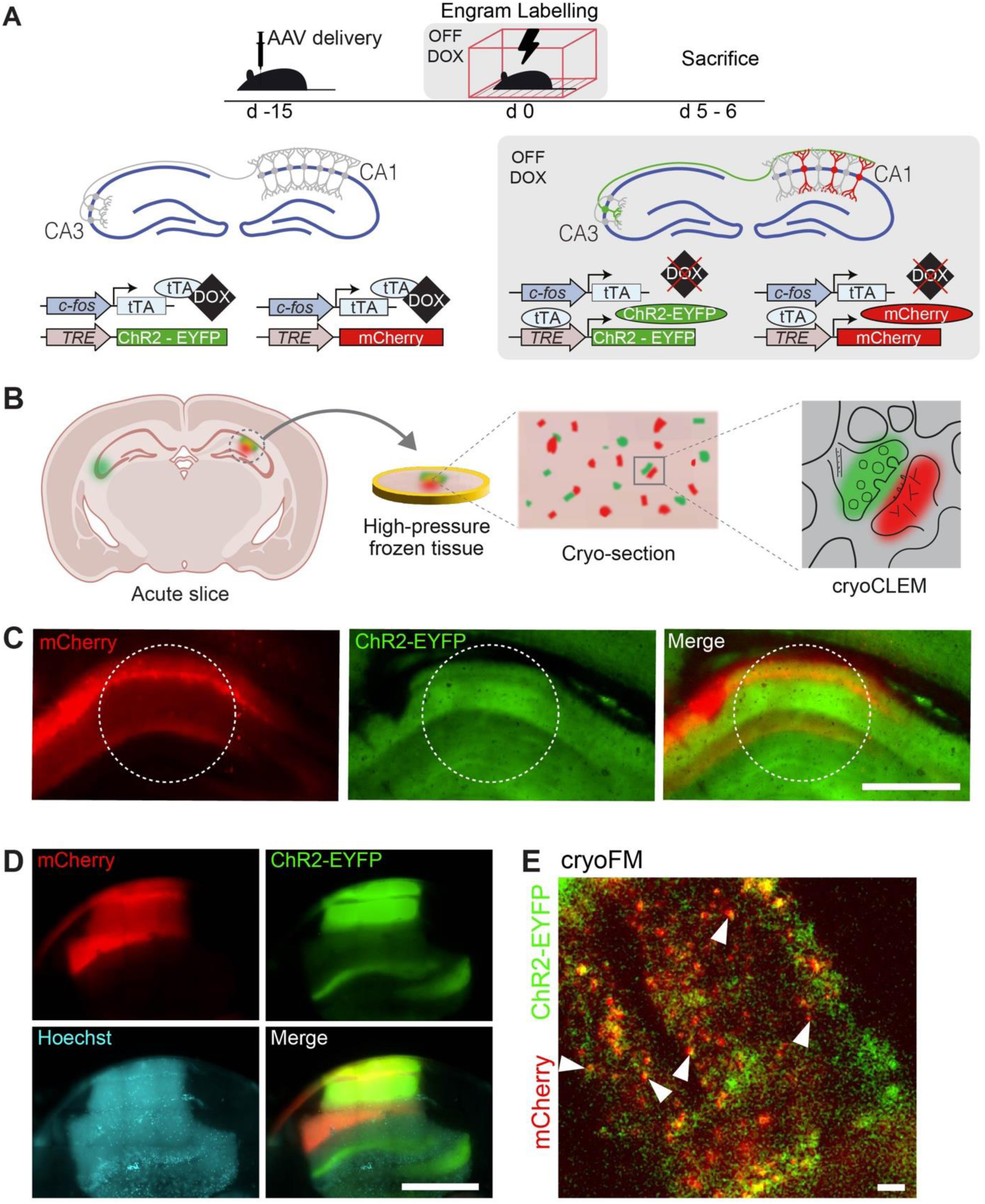
CryoCLEM targeting of engram synapses. **A.** Top and bottom, schematics showing timeline and genetic strategy to label a contextual fear memory engram, respectively. AAV-constructs were delivered into contralateral CA3 and CA1 regions to allow simultaneous labelling of pre- and post-synaptic neurons, respectively. CA3-CA1 engram was labelled by using a TRE-ChR2-EYFP vector that is activated by tTA, which expressed under the control of a c-fos promoter. In the absence of DOX (DOX OFF), engram neurons encoding an episodic memory became tagged with ChR2-EYFP (CA3, presynaptic) and mCherry (CA1, postsynaptic). Lightning symbol, foot-shock delivery; d, days **B.** Schematic depicting the detection of fluorescent engram-labelled neurons within the acute brain slices, high-pressure frozen (HPF) tissue, and cryo-sections. These fluorescent puncta were used to direct cryoET data collections of synapses. **C.** Fluorescence microscopy of engram-labelled fresh hippocampus. Green, presynaptic CA3 engram terminals (ChR2-EYFP). Red, postsynaptic CA1 engram neurons (mCherry). Blue, hoechst nuclear label. White dashed circle, area of tissue biopsy. Scale bar, 500 mm. **D.** CryoFM of high-pressure frozen (HPF) engram-labelled tissue biopsy. Colours same as C. Scale bar, 500 mm. **E.** CryoFM of engram-labelled mouse brain tissue cryo-section. Colours same as C. White arrowhead, adjacent CA3 presynaptic ChR2-YFP-positive and CA1 postsynaptic mCherry-positive puncta. Scale bar, 5 mm.

To validate that ChR2-EYFP labels the presynaptic boutons of CA3 neurons, AAV constructs encoding ChR2-EYFP under the control of a CaMKII promoter were injected into the CA3 (**Supp. Fig. 1AB**). Immunofluorescence microscopy of CA1 detected ChR2-EYFP colocalized with presynaptic markers (**Supp. Fig. 1C**), as expected^43–45^. Immunofluorescence also confirmed colocalization of mCherry and the postsynaptic density protein 95 (Psd95) (**Supp. Fig. 1D**).

To prepare engram-labelled tissue for cryoCLEM and cryoET, acute slices of fresh brain were obtained from engram-labelled mice 5-6 days after fear conditioning (**Fig. 1B**). Fluorescence microscopy of fresh acute slices showed presence of ChR2-EYFP-labelled contralateral CA3 axons and mCherry-labelled dendrites in molecular layers of CA1, corresponding to the hippocampal engram-associated to the fear memory (**Fig.1C**)^46–49^. Next, engram-labelled hippocampi were cryopreserved by high-pressure freezing (**Fig. 1D**), from which 70-150 nm tissue cryo-sections^50^ were prepared. Cryogenic fluorescence microscopy (cryoFM) of tissue cryo-sections revealed sparse labelled regions corresponding to ChR2-EYFP-tagged CA3 axons and mCherry-tagged CA1 dendrites (**Fig. 1E** and **Supp. Fig. 2**). Juxtaposition of ChR2-EYFP-labelled axons and mCherry-labelled dendrites indicated locations that could potentially contain engram to engram synapses in tissue cryo-sections (**Fig. 1E**).

We mapped the location of an engram ChR2-EYFP-tagged CA3 axon and mCherry-tagged CA1 dendrite by cryoCLEM (**Fig. 2A, B**) to direct the collection of cryoET data (tilt series with 2° tilt increments, +/-60° range, ∼110 e/Å^2^ total dose) corresponding to a 1.2 mm^2^ field of view of the tissue cryo-section with 3 Å pixel size. The reconstructed tomographic volume (**Fig. 2C**) revealed the in-tissue molecular architecture of a circuit-specific engram synapse (**Fig 2C, D, E** and **Supp. Table 1**), including macromolecular complexes and organelles. The ChR2-EYFP-containing region of the tomographic volume corresponded to a membrane bound compartment filled with presynaptic vesicles and interspersed with microtubules. In contrast, the mCherry-containing region of the tomogram marked a membrane-bound subcellular compartment containing a network of branched F-actin cytoskeleton (**Fig. 2B, C, D**). Within the cleft formed between the pre- and postsynaptic compartments, individual protein complexes were observed protruding from both the pre- and postsynaptic membranes, as well as transsynaptic complexes that tethered the pre- to the postsynaptic membrane (**Fig. 2C**). These molecular features confirm the capability of this engram to tomogram workflow to obtain the in-tissue molecular architecture of a circuit-specific engram synapse.

**Figure 2.**
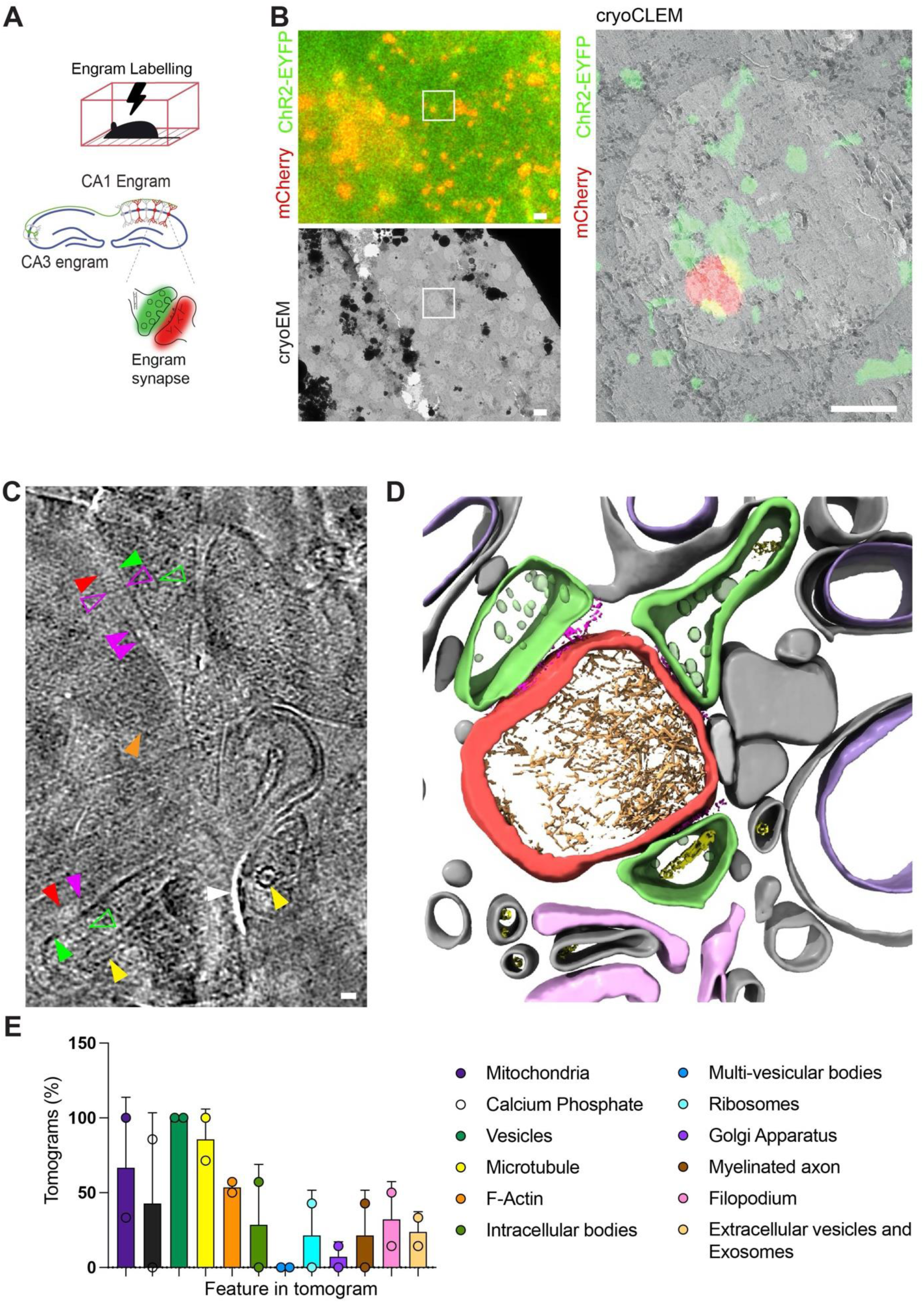
3D molecular architecture of an engram synapse by cryoET. **A.** Schematic depicting the labelling of engram cells in the CA3 and contralateral CA1 for cryoCLEM and targeted cryoET. **B.** Top left, cryoFM image of engram-labelled mouse cryo-section. Green, *TRE*-ChR2-EYFP labelled presynaptic CA3 engram neuron terminals. Red, *TRE*-Cherry labelled postsynaptic CA1 engram neuron. White box corresponds to adjacent ChR2-EYFP and mCherry puncta from which a tomographic tilt series was collected. Scale bar, 0.5 mm. Bottom left, cryoEM image of the same tissue cryosection. White box corresponds to the region from which the tomographic tilt series was collected. Scale bar, 0.5 mm. Right, aligned cryoFM and cryoEM (cryoCLEM) image of tissue cryo-section. Colours same as A. Region shown corresponds to the tilt series collected. Scale bar, 0.5 mm. **C.** Section from a tomographic slice of engram-labelled synapse. Green filled arrowhead, presynaptic membrane. Red arrowhead, postsynaptic membrane. Magenta filled arrowhead, postsynaptic membrane protein. Magenta open arrowhead, transsynaptic adhesion protein spanning the cleft. Orange arrowhead, F-Actin. Green open arrowhead, presynaptic vesicle. Yellow arrowhead, microtubule. White arrowhead, knife damage from cryo-sectioning. Scale bar, 20 nm. **D.** 3D segmentation of the reconstructed in-tissue tomogram from the location indicated in A. Green, ChR2-EYFP-positive presynaptic membrane. Red, mCherry-positive postsynaptic membrane. Dark red, cleft protein. Transparent green, presynaptic vesicle. Gold, cytoskeletal filament. Yellow, microtubules. Purple, mitochondrion, Light grey, subcellular compartments surrounding engram-labelled synapse. Pink, putative filopodia. **E.** Graph showing the prevalence of macromolecular constituents in engram synapses and surrounding compartments (n=15). Graph depicts mean per mouse ± SD.

### Quantification of engram-labelled synapses

To investigate if the composition and molecular architecture of engram synapses was conserved, we collected a dataset of fourteen in-tissue engram synapse tomograms. Adjacent engram (*TRE*-mCherry+ and *TRE*-Ch2R-EYFP+) cryoFM puncta were confirmed as engram synapses based on the correlation of EYFP to a compartment containing synaptic vesicles and of mCherry to a neighboring compartment connected via transsynaptic adhesion molecules spanning a cleft (**Supp. Table Fig. 1**). We cataloged and quantified the identifiable macromolecular and organelle constituents in these cryoET datasets (**Fig. 2E and Supp. Table 1**). Microtubules, mitochondria, and other membrane-bound organelles were present in 31%, 85%, and 50% of engram synapses, respectively (**Fig. 2E**). Ribosomes were absent from both the pre- and postsynaptic compartments of all engram synapses but were evident in non-synaptic compartments (**Supp. Table 1**).

Mitochondria were identified with and without calcium phosphate deposits^51^ (**Fig. 3A**). A large proportion of mitochondria (58%) were found within postsynaptic compartments (**Fig. 3B**)^52^. Since dendritic spines do not contain mitochondria^53,54^, it is likely that such synapses with postsynaptic mitochondria correspond to dendritic shaft synapses^55–58^.

**Figure 3.**
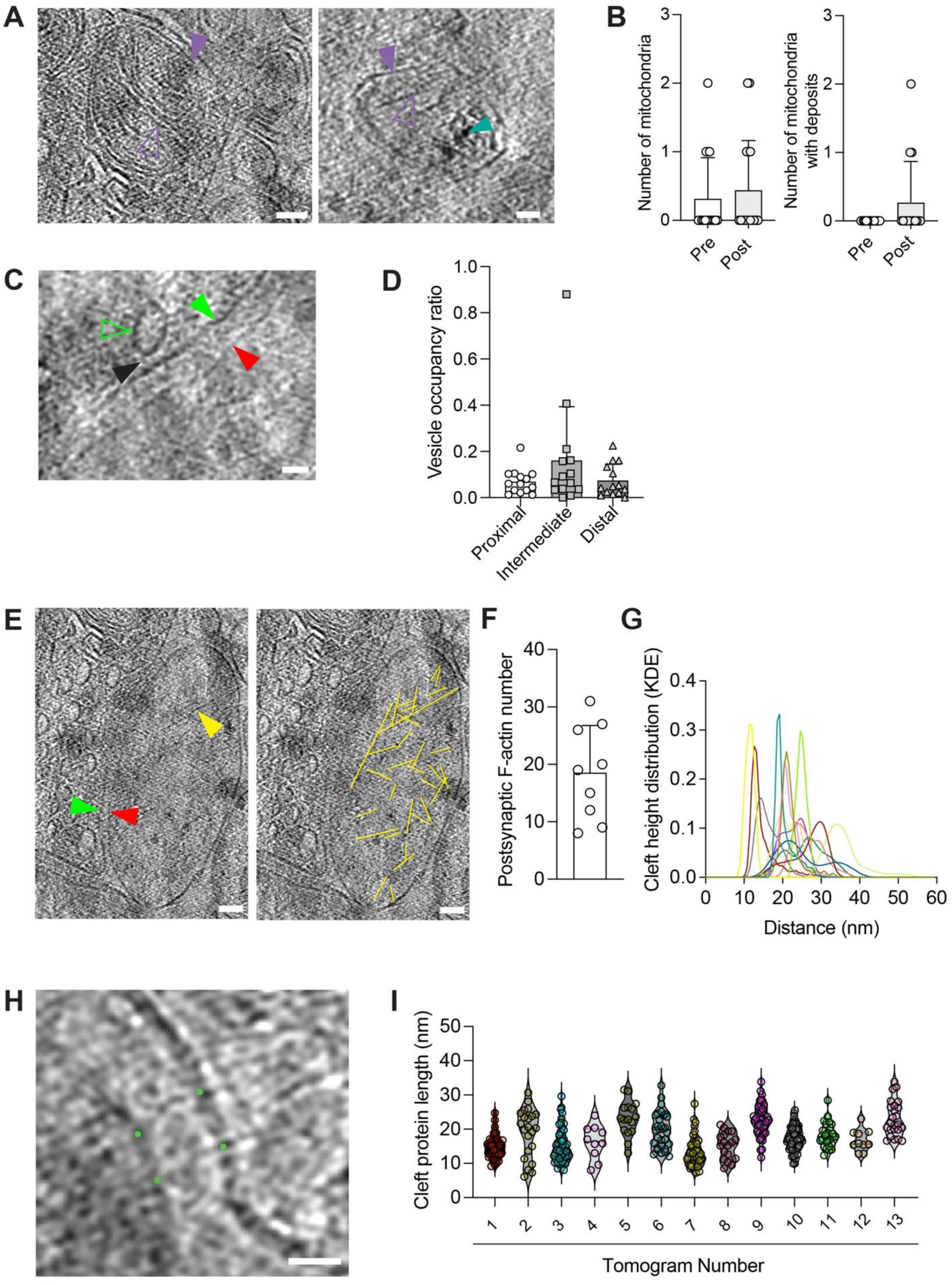
Heterogeneity of engram synapse molecular architecture. **A.** Left and right, tomographic slices of mitochondria within engram synapses without and with calcium phosphate deposits, respectively. Purple filled arrowhead, outer mitochondrial membrane. Purple outlined arrowhead, inner mitochondrial membrane forming cristae. Cyan arrowhead, calcium phosphate deposits. Scale bars, 20 nm. **B.** Left, graph showing the prevalence of mitochondria in presynaptic and postsynaptic compartments of engram synapses (mean ± SD). Right, graph showing the number of mitochondria with calcium phosphate deposits in presynaptic and postsynaptic engram-labelled compartments (mean ± SD). **C.** Tomographic slices of vesicles in an engram-labelled synapses. Green filled arrowhead, presynaptic membrane. Red arrowhead, postsynaptic membrane. Green open arrowhead presynaptic vesicle. Black arrowhead, protein tethering synaptic vesicle to presynaptic membrane. Scale bar, 20 nm. **D.** Quantification of synaptic vesicle distribution based on the ratio of the volume occupied by vesicles in proximal, intermediate and distal regions of the presynaptic compartment of engram synapses (mean per synapse ± SD). **E.** Left, tomographic slice depicting an engram-labelled synapse. Green arrowhead, presynaptic membrane. Red arrowhead, postsynaptic membrane. Yellow arrowhead, F-Actin. Right, the same tomographic volume with overlay of F-Actin segmentation indicated in yellow. Scale bar, 20 nm. **F.** The number of F-Actin filaments in engram-labelled postsynaptic compartments (mean per synapse ± SD). **G.** Distribution of cleft heights (KDE, kernel density estimation) in engram-labelled synapses. Distribution of cleft heights in each synapse shown in different colour. **H.** Tomographic slice showing measurement of the length of transsynaptic adhesion proteins. Model points (green) were placed on the pre- and post-synaptic ends of cleft adhesion proteins. The distance between these coordinate pairs was calculated to measure the length of transsynaptic adhesion proteins. Scale bar, 20 nm. **I.** Graph showing the length (mean ± SD) of cleft adhesion proteins in each engram-labelled synaptic cleft tomogram.

Presynaptic vesicles were indicative of synapses in tomograms (**Supp. Table 1**), where they play a critical role in action potential-evoked neurotransmission^59^. Synaptic vesicles with 33 ± 1.22 nm (mean ± SD) diameter were variously distributed throughout presynaptic compartments, as expected^34,38,60–62^. The distribution of synaptic vesicles could give structural insight into presynaptic mechanism contributing to the synaptic strength of each synapse because only vesicles proximal to the active zone are thought capable of mediating fast synaptic transmission^33,61–63^. Synaptic vesicle distribution was assessed by measuring the total volume occupied by proximal (0-45 nm from presynaptic membrane, **Fig. 3C**), medial (45 - 75 nm from presynaptic membrane) and distal (> 75nm from presynaptic membrane) synaptic vesicles. There was no significant difference between the occupancy at these locations (proximal versus intermediate and distal vesicle populations) (**Fig. 3D**), which was comparable to that previously reported in primary neurons^33^.

F-actin cytoskeleton in postsynaptic compartments plays a central role in structural remodelling during synaptic plasticity^6,64–67^, particularly at dendritic spines^68^. We mapped cytoskeletal networks in each postsynaptic compartment by segmenting and analyzing filaments in each engram synapse^69^ (**Fig. 3E, F**). These data showed F-actin formed branched networks, with a 3-fold variation of copy number, and were a conserved constituent of engram synapses^34^.

The strength of synaptic transmission is dependent upon multiple factors, including geometric parameters of the synaptic cleft^70^. We measured the cleft height (nearest-neighbour distance between the pre- and postsynaptic membrane), revealing the mean cleft height of each engram synapse tomogram varied from 12 to 35 nm (**Fig. 3G**). This range was larger than previously reported in analysis of 2D EM images of chemically fixed and heavy metal-stained synapses^70,71^. However, cleft height distribution was comparable to that reported for cryosections of PSD95-EGFP-labelled glutamatergic synapses^34^. Some clefts had a bimodal cleft height distribution, inferring that there were distinct subregions of synaptic cleft. We measured the height of cleft adhesion proteins in the synaptic cleft (**Fig. 3H, I**), demonstrating a similar distribution to the cleft height, suggesting adhesion proteins modulate the local cleft height of engram synapses.

To analyze further the distribution of proteins in the synaptic cleft we performed subtomogram average analysis^72,73^. We used an unbiased approach to pick proteins by over-sampling the pre- and postsynaptic membrane within the cleft. Alignment and PCA analysis classified those subvolumes containing individual proteins into six classes each containing 21-70 subvolumes and a copy number of up to 29 membrane proteins per class per synapse tomogram **(Supp. Fig2A, B)**. These low-resolution class averages were consistent with the presence of cleft resident proteins but were of insufficient resolution to definitively identify constituents (**Supp. Fig. 2C-H**), for which a copy number of at least ∼4000 is necessary to resolve the protein fold that identifies specific individual proteins within tissue cryo-sections by cryoET^74^.

### Analysis of non-synaptic subcellular compartments

Non-synaptic regions adjacent to engram synapses (*TRE*-ChR2-EYFP+ or *TRE*-mCherry+ cryoCLEM) were also captured within our in-tissue cryoET dataset (**Supp. Table 1**). Myelinated axons accounted for 14% non-synaptic engram-labelled subcellular compartments. mCherry-labelled axons were absent, consistent with the CA3 (presynaptic) targeting of *TRE*-ChR2-EYFP expression (**Supp. Fig. 3A**). Filo- or lamellipodia accounted for 16% non-synaptic *TRE*-ChR2- EYFP+ subcellular compartments, with 18-46 nm average diameters (**Supp. Fig. 3B**). These subcellular structures could correspond to cellular intermediates of synaptogenesis^75–77^. The remaining 70% of non-synaptic *TRE*-ChR2-EYFP+ subcellular compartments (**Supp. Table 1**) likely corresponded to unmyelinated axonal processes^78^. Non-synaptic TRE-mCherry+ membrane-bound subcellular compartments likely corresponded to somato-dendritic regions of CA1 neurons (**Supp. Fig. 3C**).

We also analyzed unlabeled (*TRE*-ChR2-EYFP- and *TRE*-mCherry- cryoCLEM) subcellular compartments surrounding each synapse that provided in-tissue contextual information in the form of multiple membrane-enclosed subcellular compartments (17 to 627 nm diameter) (**Supp. Table 1**). The cellular origin, either neuronal or non-neuronal, for most of these subcellular compartments could not be definitively determined. Nonetheless, 15% of engram synapse tomographic volumes contained unlabeled filo- or lamellipodia (**Supp. Fig. 3A, *right***), 33% tomographic volumes containing synapses also contained unlabeled myelinated axons (**Supp. Fig. 3B, *right***), and 19% contained extracellular vesicles (**Supp. Fig. 3D and Supp. Table 1**)^79^. The proximity of unlabeled to engram-labelled subcellular compartments is consistent with the sparse distribution of neurons that form engram circuits^2,16,80^ and that engram circuits are not spatially segregated from other neuronal circuits^2,16,80^.

### Cell-specific cryoCLEM-guided cryoET

To test further the general application of targeting genetically labelling subpopulations of neurons for cryoET, we collected an additional nine synapse tomograms formed from engram-labelled ChR2-EYFP-positive CA3 presynaptic terminals and unlabeled contralateral CA1 postsynaptic terminals **(Fig. 4A and Supp. Table 1).** ChR2-EYFP marked a presynaptic compartment containing numerous presynaptic vesicles, confirming the fidelity of cryoCLEM-guided cryoET to locate synapses with a single engram cell fluorescent label. To test if this cryoCLEM-guided cryoET could identify synapses within larger labeled neuronal ensembles, an additional three synapse tomograms were also collected from mice receiving pan-neuronal presynaptic labelling in CA3 (with *CamKII*-ChR2-EYFP AAV) and *TRE*-mCherry AAV for postsynaptic labelling in CA1. **Fig. 4B**, **Supp. Fig. 1A**, and **Supp. Table 1**). Pan-neuronal ChR2-EYFP and mCherry mapped to cellular compartments containing presynaptic vesicles and a network of branched F-actin, respectively. These data further demonstrate the broad versatility of this workflow to obtain molecular resolution tomographic maps of specific synapses within neuron ensembles in the mammalian brain.

**Figure 4.**
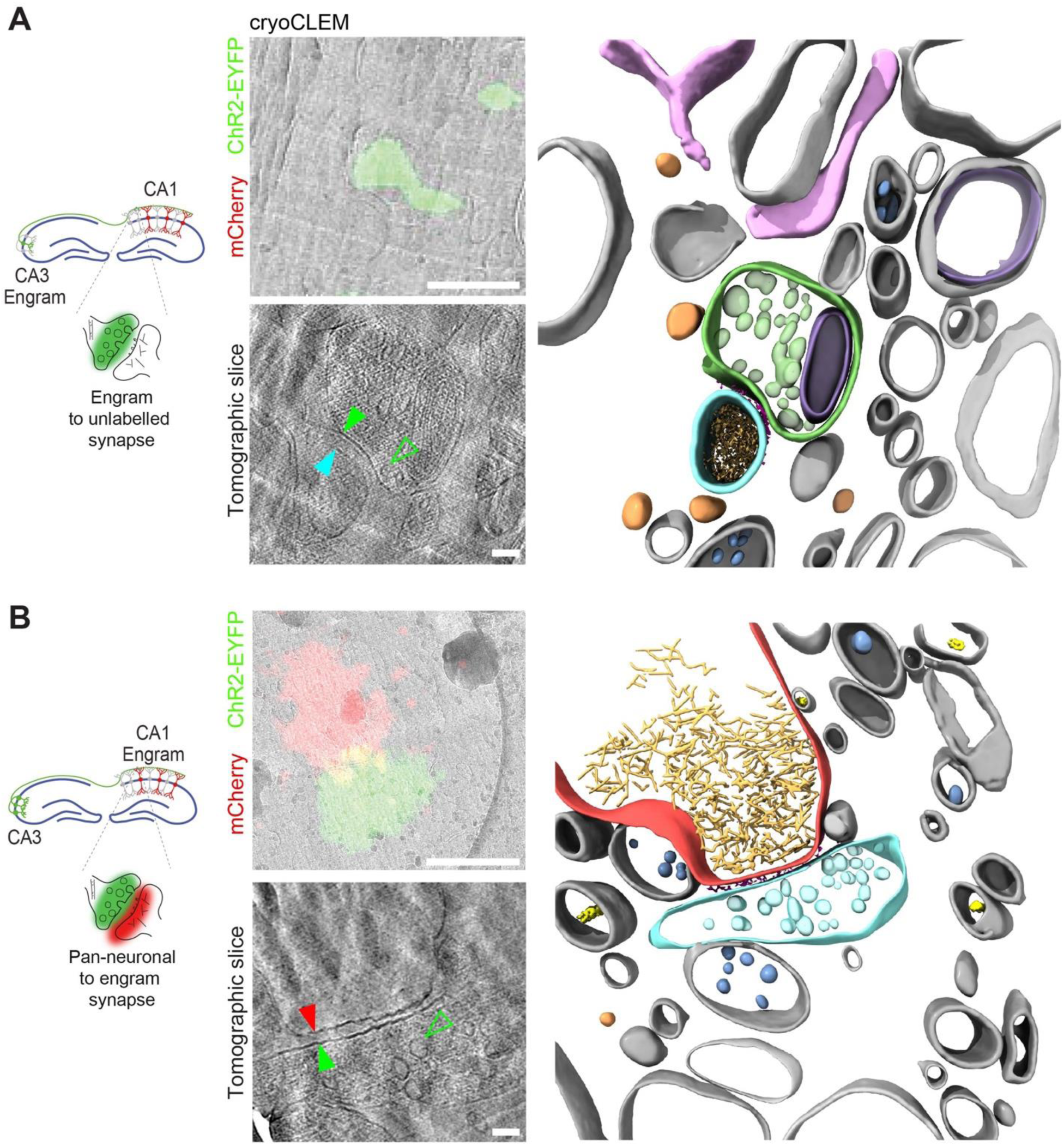
CryoCLEM-guided cryoET of neuronal ensembles. **A.** CryoCLEM-targeted cryoET of a synapse between an engram-labelled cell and an unlabelled cell in a tissue cryo-section. Left, schematic depicting the labelling of engram cells in the CA3 and contralateral CA1. To survey the surrounding synapses, tomograms were collected of synapses between engram-labelled CA3 neurons and unlabelled CA1 neurons. Middle top cryoCLEM. Green, *TRE*-ChR2-EYFP labelled presynaptic CA3 engram neuron terminal. Scale bar, 0.5 mm. Middle bottom tomographic slice of a synapse between an engram-labelled cell and a neuron outside of the labelled engram circuit. Green filled arrowhead, engram-labelled presynaptic membrane. Cyan arrowhead, unlabelled postsynaptic membrane. Green open arrowhead, presynaptic vesicle. Scale bar, 20 nm. Right, segmentation of tomogram shown bottom left. Green, ChR2-EYFP-positive presynaptic membrane; Cyan, unlabelled postsynaptic membrane; Dark red, cleft proteins. Transparent green, presynaptic vesicle. Gold, cytoskeletal filament. Yellow, microtubule. Dark grey, non-synaptic vesicle. Light red, extracellular vesicle. Pink, putative filopodia. Purple, mitochondrion. Light grey, subcellular compartments surrounding engram-labelled synapse. **B.** CryoCLEM-targeted cryoET of a synapse from a *CamKII*-ChR2-EYFP/TRE-mCherry-labelled mouse cryo-section. Left, schematic depicting the labelling of CA3 cells with a pan-neuronal CaMKII marker and engram cells in the contralateral CA1. Tomograms of synapses with CA1 engram cells were collected. Middle top, cryoCLEM. Green, *CamKII*-ChR2-EYFP labelled presynaptic CA3 neuronal terminal. Red, *TRE*-Cherry engram-labelled postsynaptic CA1 neuron. Scale bar, 0.5 mm. Middle bottom, tomographic slice of a synapse between an engram-labelled postsynaptic neuron and a pan-neuronal (*CamKII*)-labelled presynaptic neuron. Green filled arrowhead, pan-neuronal presynaptic membrane. Red arrowhead, engram postsynaptic membrane. Green outlined arrowhead, presynaptic vesicle. Scale bar, 20 nm. Right, segmentation of tomogram in bottom left. Colours same as in A.

## Discussion

Here, we developed and applied a novel workflow bridging animal behaviour to proteins within neuronal circuits, unveiling the 3D molecular architecture of synapses that underlie engram connectivity. We described and quantified the heterogeneity of these synapses, demonstrating that diversity exists within engram contacts with regards to organelles, and macromolecular complexes. This workflow provides a foundation for further interrogating engram cell connectivity in fresh tissue at the subcellular and molecular level by in-tissue cryoET^34,74^.

Earlier studies of synapse structure cryopreserved neuronal cultures and brain tissues to analyze synapses by cryoET^31,34–36,39,81,82^ and cryoCLEM labelling has previously been applied to cellular and tissue cryoET^34,35,39^. The recent application of room temperature, volumetric EM of resin-embedded tissue to the study of engram cells allowed the characterization of the cellular morphology and the connectivity of CA1 engram neurons one week after memory acquisition^83^. This elegant, complementary study discovered CA1 engram neurons contained a higher fraction of multi-synaptic boutons compared to CA1 neurons in control mice. However, engram synapse identification requires pre- and postsynaptic identification of engram circuit components. This study was therefore unable to distinguish engram from non-engram synapses per se. While resin-embedded volumetric EM interrogates overall changes in the extent of connectivity, cryoCLEM-targeted cryoET additionally gives insight into the native 3D macromolecular architecture of engram versus non-engram synapses.

The 3D architecture, including organelle and macromolecular composition, of engram synapses were highly variable, likely reflecting known diversity of synapse types even within the same brain region^84,85^. We also observed variable cleft height within and between synapses^34^. This heterogeneity could be explained, at least in part, by the fact that both excitatory and inhibitory neurons were potentially labelled during memory encoding. Indeed, synapse tomograms included those both on dendrites and within dendritic spines. Future studies could further dissect engram synapses by incorporating additional cell type-specific genetic reporters or targeting markers of structural plasticity^67^, as well as combining volumetric cryoFIB-SEM imaging with cryoET of FIB-milled lamella^86^.

Learning and memory-related plasticity mechanisms have been reported to affect specific cellular protein constituents of neuronal circuits at different times^67,83,87,88^. Developing engram circuit labelling methods to acquire in-tissue cryoET dataset at early versus late time-points following memory encoding could determine if structural differences transiently exist or persist at time points relevant to plasticity processes. This could resolve specific changes at engram synapses relating to different phases of memory encoding, such as recall, consolidation and storage, and relating to different brain regions^5,21,80,89^. Moreover, the combination of our workflow with both natural or artificial optogenetic memory recalls could enable the comparison of engram synapses between active and inactive memory circuits^90–93^.

The ‘engram to tomogram’ workflow reported here provides a first demonstration that sparse engram synapses can be identified and visualized at macromolecular resolution within the mouse brain. Larger cryoET datasets of engram synapses could allow direct determination of subnanometer resolution in-tissue protein structures by subtomogram averaging^74^. Additionally, the development of in-tissue cryoET methods to collect tomograms from synapses spanning several serial cryo-sections could enable a more complete structural interrogation of each engram synapse^82^. Application of these targeted cryoET approaches will necessarily expand the engram field to relate behavioural and electrophysiological properties ^2,17,94^ to their structural counterparts and gain a deeper understanding of plasticity mechanisms that mediate different stages of memory encoding. More broadly, defining the synaptic and structural plasticity mechanisms behind engram formation and function is key to understanding how the brain computes information to adapt to changing environments^95^ and could also provide insight into the molecular changes to engram synapses associated with development, aging, stress, trauma, and models of neuropsychiatric and neurodegenerative disorders^96–106^.

## Acknowledgements

C.L. was funded by a BBSRC White Rose DTP PhD studentship (BB/M011151/1). T.R. and C.O.S. were funded by the European Research Council (715968), Science Foundation Ireland (15/YI/3187), the National Institute of Health (1R01NS121316) and the Irish Research Council (GOID/2019-812). R.F. was funded by an Academy of Medical Sciences Springboard Award (SBF005/1046), a UKRI Future Leader Fellowship (MR/V022644/1) and a University of Leeds Academic Fellowship. The Astbury Biostructure Laboratory Titan Krios microscopes were funded by the University of Leeds and Wellcome (108466/Z/15/Z & 221524/Z/20/Z). The Leica EM ICE, UC7 ultra/cryo-ultramicrotome and cryoCLEM systems were funded by Wellcome (208395/Z/17/Z). We thank Lydia Marks and Mariia Yurova for technical and administrative support, and all the past and present members of the Ryan Lab for support and discussion. We also thank Kawar Abbas, Andrew Horner, and Melanie Reay for technical support. We thank Pasquale Pelliccia, Jon Whitwell, Wallace Tudeme, Kerry Higgins, and Martin Foster for computational support. We thank Rebecca Thompson, Emma Hesketh, Louie Aspinall, Joshua White, Yehuda Halfon and Oksana Degtjarik for help maintaining and setting up the Astbury Biostructure Laboratory (ABSL) cryoEM facility, including high-pressure freezing, cryoCLEM and Titan Krios microscopes. We thank Ruth Hughes and Sally Boxall for maintaining and training at the bioimaging facility, including the confocal and wide-field fluorescence microscopes.

## Author contributions

C.O.S. conducted surgeries, behavioral and engram labelling experiments. C.L performed brain dissection, acute brain slice preparation, and high-pressure freezing, C.L. and T.O. performed tissue cryo-sectioning. C.L. performed immunofluorescence, cryoFM and cryoET data collection and processing, and computational image analysis. T.J.R., R.F., C.O.S. and C.L. interpreted the data. C.L. and C.O.S. wrote the original draft. All authors reviewed and edited the final manuscript. C.O.S., T.J.R. and R.F. conceived the scientific design and supervised the project.

## Methods

### Animals

The experimental subjects employed were male C57Bl6/J mice (Charles River), aged 7 to 14 weeks. The animal room was maintained at a constant temperature of 22°C, and a 12-hour light/dark cycle was established, with all experimental steps conducted during the light phase. Mice were housed in groups of five, in cages equipped with a tunnel, with free access to food and water. The care and behavioural experiments involving the mice were approved by the Animal Ethics Committee of Trinity College Dublin, University of Leeds Animal Welfare and Ethics Committee. Experiments were conducted in accordance with the Health Products Regulatory Authority of Ireland, European Directive 2010/63/EU, and the UK Animal Scientific Procedures Act.

### Stereotactic surgery and engram labelling

Stereotactic surgery was performed under stereotactic guidance using standard mouse stereotactic frames. Mice were anaesthetized using 500 mg/kg avertin (Sigma). During the procedure, a bilateral craniotomy using a 0.5 mm diameter drill was carried out and viral cocktails were injected through a metal needle attached to a 10 µL Hamilton microsyringe (701LT; Hamilton) and an automated microsyringe pump (WPI). To label engram cells, *cfos-tTA*, *TRE-mCherry* and *TRE-ChR2-EYFP* AAV plasmids were used (gift from S.Tonegawa). For pan-neuronal labelling, *TRE-ChR2-EYFP* was substituted for *CamKII-ChR2-EYFP* (Addgene, 26969). Plasmids were AAV9 serotyped and packed into viral particles by Vigene Bioscience (Maryland, USA). The needle was placed on appropriate stereotactic coordinates and remained for 5 min before the injection commenced. The coordinates used were: CA1 (-2.0 mm AP, +1.3 mm ML, – 1.4 mm DV) and contralateral CA3 (-2.0 mm AP, -2.5 mm ML, –2.3 mm DV); 250 nl or 200 nl (for CA1 or CA3 respectively) of cocktail virus were injected at 60 nl/min speed. After the injection, the needle stayed for ten additional minutes before it was carefully withdrawn. The incision was closed with sutures. Mice were given 1.5 mg/kg metacam (Meloxicam) as an analgesic once per day for two days after surgery. Once returned to the home cage, animal health was assessed every two-three days. Mice were allowed to recover for ten days prior to engram labelling. Mice were fed 40 mg kg-1 doxycycline (DOX) for at least a week before surgery. Animals were individually handled for three min each day for three days immediately before the engram labelling. On the fourth day, the DOX diet was substituted for regular diet. After 36 hours, animals were subjected to Contextual Fear Conditioning. Animals were transported to an experimental room where they were allowed to explore a context for 3 minutes, followed by 3 successive 0.75 mA shocks of 2 s duration spaced by one minute. Contextual cues used were a triangular shape inset and Benzhaldehyde 0.25% (Med Associates Contextual Chambers). Immediately after CFC, animals were put back on DOX diet. Four adult male mice were used for tomography data collection.

### Immunohistochemistry

Mice were sacrificed 8 days after engram labelling and brains were collected. Whole brains from 3 adult male mice were flash-frozen in liquid nitrogen. Brains were thawed, flash-frozen in OCT and mounted in a cryostat (Leica). At -18°C, 14 μm thick brain slices were cut and attached to glass slides. Slices were fixed in ice-cold methanol for 7 minutes and were stained with antibodies against Synapsin-1 (Invitrogen cat #A-6442, 1:200) or PSD-95 (Neuromab, cat #75-028, 1:200) at 4°C overnight. The next day, slides were washed in PBS. A secondary antibody was applied (ThermoFisher Anti-Rabbit-AF 568, 1:1000, or Anti-IgG2a-AF 594, 1:1000) for 2 hours at room temperature. Tissue sections were washed in PBS and mounted in Vectashield mountant with DAPI (Vector Laboratories, Burlingame, CA).

All images were captured using a confocal laser scanning microscope (Zeiss LSM 700) utilising a 10x air objective lens (0.45 numerical aperture) and 63x oil objective lens (1.2 numerical aperture) with frame size 1024x1024 pixels. Samples were imaged using 405 nm, 488 nm and 561 nm lasers. Colocalization was measured using the Coloc2 package in Fiji and the Pearson’s correlation value was plotted.

### Preparation of acute slices

Mice were 6 days after engram labelling administered intraperitoneal injection of pentobarbital (100 mg/kg) and intracardial perfusion of room temperature NMDG cutting buffer (23.25 mM NMDG, 0.625 mM KCl, 0.3 mM NaH2PO4, 7.5 mM NaHCO3, 5 mM HEPES, 6.25 mM C₆H₁₂O₆, 1.25 mM C6H7O6Na, 0.5 mM CH4N2S, 0.75 mM C3H3NaO3, 2.5 mM MgSO4.7H2O, 0.125 mM CaCl2.2H2O; pH 7.2-7.4; 304-310 mOsm; adapted from^107^. Coronal sections 100 µm thick were prepared using a Leica vibratome (0.26mm/s) in ice-cold NMDG cutting buffer. For cryoCLEM and cryoET experiments, the slices were recovered in room temperature hACSF buffer (120 mM NaCl2, 5 mM KCl, 1.2 mM MgCl2.6H2O, 2 mM CaCl2.2H2O, 25 mM HEPES, 30 mM C₆H₁₂O₆; pH 7.2-7.4^108^.

### High-pressure freezing

Acute slices were incubated in 1 µM Hoechst 33342 (ThermoFisher, cat #62249) in NMDG cutting buffer at room temperature to label the cell bodies for subsequent identification of the granular layer. Acute slices were imaged on an EVOS Auto2 microscope with a 4x air objective (0.13 Ph LWD) and equipped with DAPI (Ex357/44, Em 447/60), GFP (Ex470/22, Em 510/42) and RFP (Ex531/40, Em 593/40) filter cubes (Invitrogen). Biopsy punches were taken from regions of interest using a 1.2 mm diameter tissue puncher and were incubated in cryoprotectant (20% dextran in NMDG cutting buffer^38^ for 30 mins at room temperature. 100 μm deep wells inside 3 mm diameter A-type gold carriers were filled with cryoprotectant and tissue. The A- and lipid-coated B- type carriers were loaded into the cartridge of the Leica EM ICE and were high-pressure frozen. Carriers were stored in liquid nitrogen.

### Cryo-ultramicrotomy

High-pressure frozen carriers were mounted into the specimen holder of the Leica EM FC7 for cryo-sectioning. Carriers were trimmed with a diamond knife (Diatome, Trim20) and 100-190 nm thick sections were cut at -160 °C using a cutting knife (Diatome, cryo-immuo). Sections were pulled into ribbons with a gold eyelash using a micro-manipulator^109^. Ribbons were transferred onto glow-discharged 200 mesh 3.5/1 Cu grids (Quantifoil Micro Tools, Jena, Germany).

### Cryogenic fluorescence microscopy (cryoFM)

Cryo-sections attached to EM grids were imaged using a cryogenic fluorescence microscope (Leica Thunder) with a HC PL APO 50x/0.9 NA cryo-objective, Orca Flash 4.0 V2 sCMOS camera (Hamamatsu Photonics), a Solar Light Engine (Lumencor) and GFP (450-490 nm excitation; 500-550 nm emission), EYFP-ET (500/20 excitation, 535/30 emission), Rhodamine (541-551 nm excitation; 565-605 nm emission) and CY5 (608-648 nm excitation; 692-740 nm emission) filter sets. An additional zoom factor of 5 was applied and images with frame size 2048 x 2048 pixels were obtained. Tilescans of carriers and grids were acquired using the LASX navigator. Z-stacks of areas of interest were acquired with 30 % intensity and 0.2 s exposure time. Images were processed using Fiji ImageJ.

### Cryogenic correlated light and electron microscopy and cryo-electron tomography

EM Grid squares were selected for electron microscopy based on their fluorescent puncta and grid orientation using ThermoFisher MAPS 3.0 software. Tomograms were collected using an FEI Titan Krios and autoloader (camera: Selectris energy filtered Falcon 4) with 3 Å pixel size. High precision correlation was achieved using MatLab scripts^110^ (Schorb and Briggs, 2014). Tomograms were reconstructed from their respective tilt series from +60° to –60° in increments of 2° using a dose symmetric tilt-shift using Tomo 5.8 software. Tilt series were collected with 2 s exposure at a dose of 0.9 Å^2^/s and between -5 to -8 µm defocus, resulting in a total dose of ∼109 electrons and pixel size of 3 Å. Dose fractions were aligned, and tomograms were reconstructed using patch tracking in IMOD software^111^. Tomograms were deconvolved in IsoNet^112^.

### Subtomogram averaging

To subtomogram average synaptic cleft proteins, membrane oversampling models were generated in Dynamo^113^. Six iterations of averaging were performed using a 20x20x10 box size and a cylindrical mask in PEET^73^. Classification into six classes was achieved with PCA analysis in PEET. ChimeraX 1.6^114^ was used to visualize classes in 3D.

### Data Handling and Statistical Analysis

Graphs were produced using GraphPad Prism software. Fluorescence images were analysed in Fiji ImageJ. Vesicle analysis was carried out using a combination of IMOD software^111^ and MatLab scripts. Cleft height was measured using Dynamo and MatLab scripts^34^. Cytoskeletal filament number was measured using an ultrastructural analysis toolkit^69^. Segmentations were prepared using a combination of IMOD, Dynamo and ChimeraX 1.6 software.

### Identification of constituents within tomographic volumes

Constituents were identified (**Supp. Table 1**) based upon the following criteria. Intracellular versus extracellular regions were distinguished by the higher density of cellular cytoplasm and characteristic intracellular organelles. Myelinated axons were defined as central compartment containing membrane-bound organelles and surrounded by rings of membrane lipid bilayer^115^. Filopodia/lamellapodia were defined as <60 nm diameter membrane-bound protrusions with a closed tip within the tomographic volume^116^. Extracellular vesicles were defined as closed vesicles situated within interstitial regions of the tomographic volume^79^. Mitochondria were defined as double membrane-bound organelle with inner membrane forming folded cristae. Calcium phosphate deposits were identified as electron-dense granules evident within mitochondria ^51^. Multi-vesicular bodies were identified as a group of two or more vesicles found encompassed by a larger membrane within an intracellular space. F-actin was defined as ∼7 nm diameter filaments composed of a helical arrangement of globular subunits^117^. Microtubules were defined as 25 nm diameter tubes composed of 13 tubulin subunits. Golgi apparatus was identified based on the presence of a stack of membranes in an intracellular compartment. Membrane organelles of unknown identity (**Supp. Fig 3D**) were defined as >80 nm diameter intracellular membrane-bound compartments.

## Supplemental Figures and Table

**Supplementary Figure 1.**
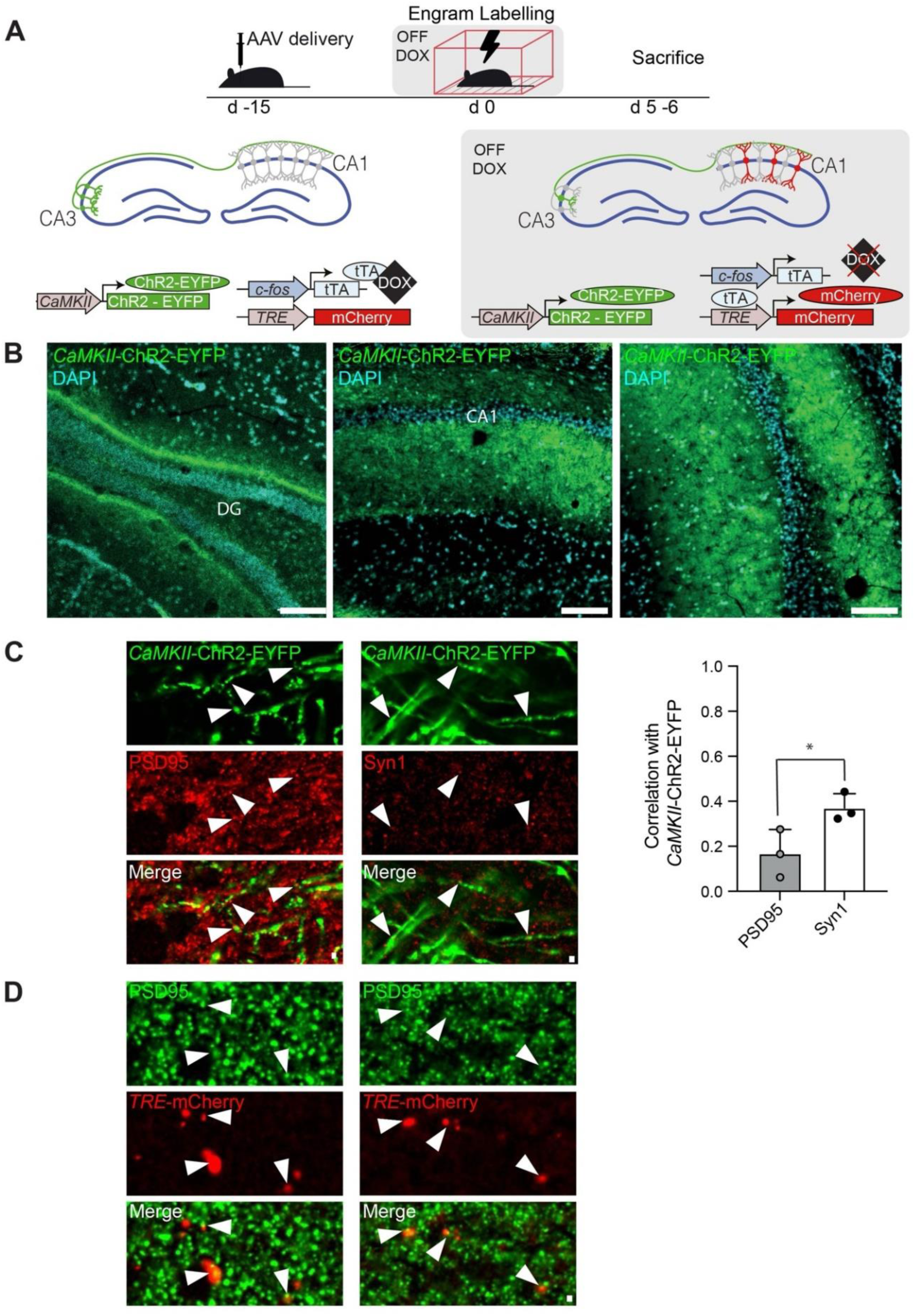
Confirmation of engram-labelled synapses in the hippocampus. **A.** Constructs were delivered intro contralateral CA3 and CA1 hippocampal regions to allow simultaneous labelling of presynaptic and postsynaptic components respectively. Projections into CA1 were labelled by contralateral CA3 injection of *CaMKII-ChR2-EYFP* AAV, whereas engram-specific labelling was achieved with the doxycycline (DOX)-controled AAV cocktail *cFos-tTA*; *TRE-mCherry*. In the absence of DOX, engram neurons encoding for an episodic memory (contextual fear conditioning) became tagged with mCherry. Lightning symbol represents foot-shock delivery. **B.** Cryostat sections of *CaMKII-ChR2-EYFP* AAV-injected brain tissue from the contralateral side, indicating the DG, CA1 and CA3 regions. Scale bars, 100 µm. **C.** Left, cryostat sections of *CaMKII-ChR2-EYFP* AAV-injected brain tissue from the contralateral CA1 region labelled with antibodies against PSD95 (left) and Synapsin-1 (right). Scale bar, 1 µm. Right, Colocalisation analysis (coloc2, see methods) shown as Pearson’s correlation coefficient, indicating negligible correlation of CaMKII-ChR2-EYFP and PSD95 and low-moderate correlation of *CaMKII*-ChR2-EYFP and synapsin-1-alexa-594. * p<0.05 via two-tailed Student’s t-test. Data presented as mean per acute slice ± SEM, with at least 3 images taken per slice. N= 3 mice. **D.** Cryostat sections of *TRE-mCherry* AAV-injected brain tissue from the CA1 region labelled with an antibody against PSD95. Scale bar, 1 µm.

**Supplementary Figure 2.**
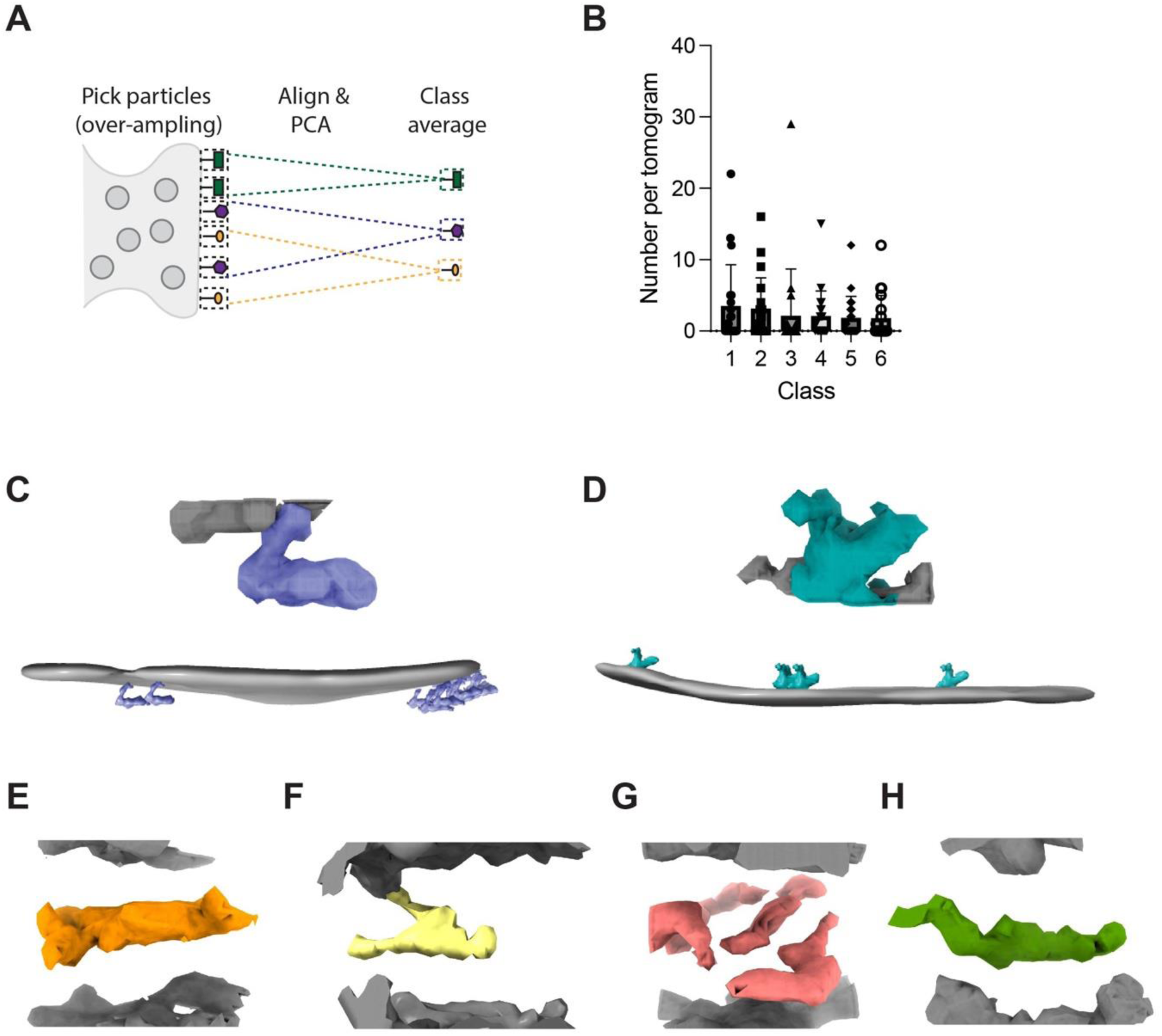
Subtomogram averaging of molecular constituents in the synaptic clefts of CA1 region synapses. **A.** Schematic representation of subtomogram averaging, picking subvolumes by membrane oversampling, followed by alignment, and PCA/*k*-means classification in PEET (see methods) to generate class averages of synaptic cleft constituents. **B.** Copy number of each class average per tomogram (mean ± SD). **C.** Top, tomographic density of class average ‘3’. Purple and grey tomographic density correspond to extracellular region of protein and membrane bilayer of engram synapse, respectively. Bottom, raw tomographic density of engram synapse showing the distribution of class average ‘3’ in cleft membrane. **D.** Top, tomographic density of class average ‘2’. Cyan and grey tomographic density correspond to extracellular region of protein and postsynaptic membrane bilayer of engram synapse, respectively. Bottom, raw tomographic density of engram synapse showing the distribution of class average ‘2’ in cleft membrane. **E.** Tomographic density of class average ‘1’. Orange and grey tomographic density correspond to extracellular region of protein and lipid bilayers, respectively. **F.** Same as **E** but for class ‘4’ in yellow. **G.** Same as **E** but for class ‘5’ in red. **H.** Same as **E** but for class ‘6’ in green.

**Supplementary Figure 3.**
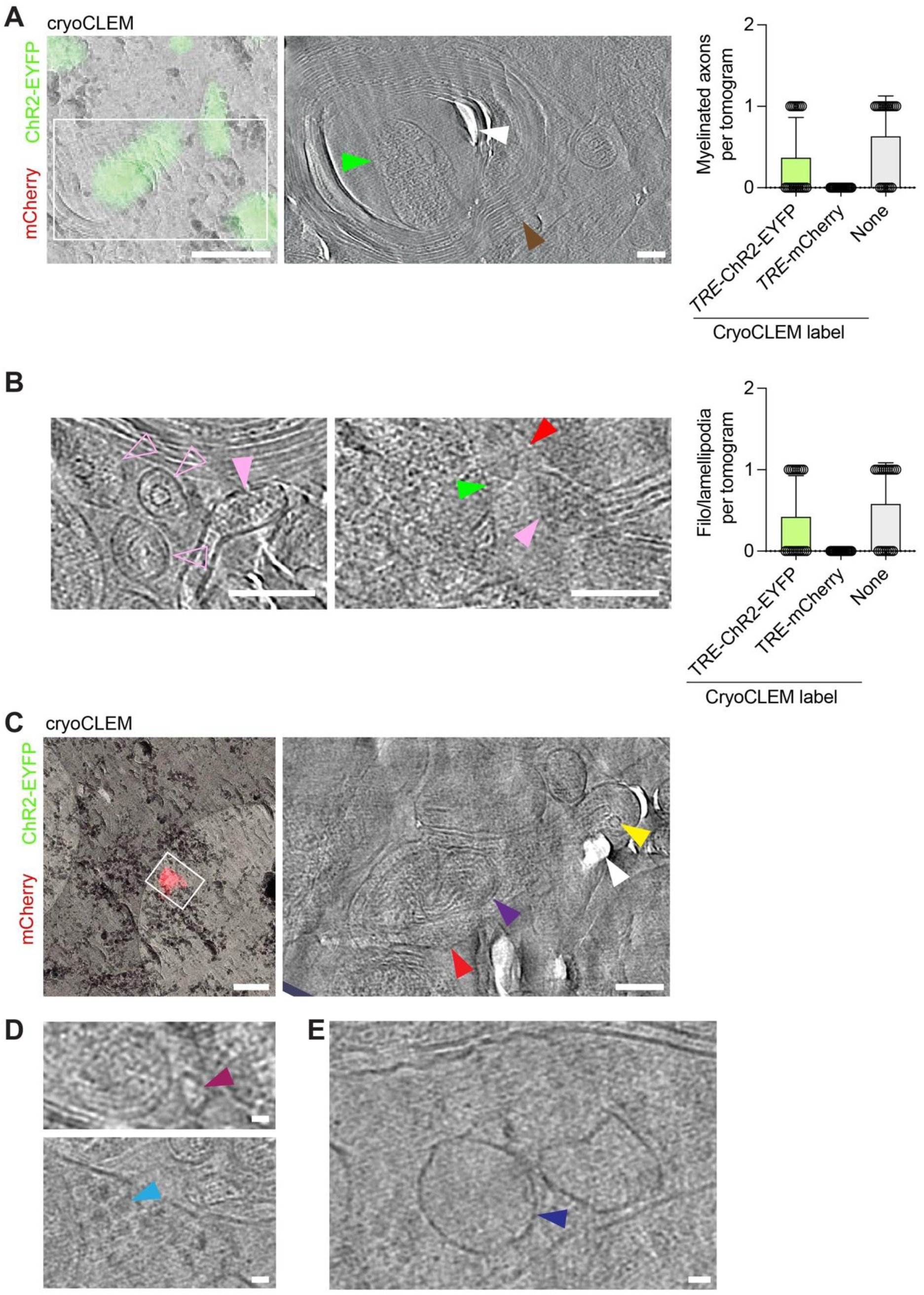
cryoET of non-synaptic constituents. **A.** CryoCLEM-guided cryoET of myelinated axons. Left, cryoCLEM. Green, ChR2-EYFP. White box, region corresponding to tomographic slice shown in middle panel. Scale bar, 500 nm. Middle, tomographic slice showing ChR2-EYFP+ myelinated axon. Green arrowhead, membrane-bound organelle. Brown arrowhead, myelin. White arrowhead, cryo-sectioning knife. Scale bar, 50 nm. Right, prevalence of ChR2-EYFP+, mCherry+ and unlabelled (ChR2-EYFP-/mCherry-) myelinated axons in cryoET dataset (mean <u>±</u> SD) (**Supp. Table 1**). **B.** CryoCLEM-guided cryoET of filo/lamellipodia. Left and middle, tomographic slice of ChR2- EYFP-/mCherry- filo/lamellapodia. Pink arrowhead, putative filo/lamellapodia. Open pink arrowhead, putative filopodia containing microtubules. Green arrowhead, presynaptic membrane. Red arrowhead, postsynaptic membrane. Scale bar,100 nm. Right, prevalence of ChR2-EYFP+, mCherry+ and unlabelled (ChR2-EYFP-/mCherry-) filo/lamellapodia in cryoET dataset (mean <u>±</u> SD) (**Supp. Table 1**). **C.** CryoCLEM-guided cryoET of non-synaptic compartment of CA1 labelled engram cell (mCherry+). Left, cryoCLEM. Green, ChR2-EYFP, Red, mCherry. White box, region corresponding to tomographic slice shown in middle panel. Scale bar, 500 nm. Right, tomographic slice showing non-synaptic subcellular compartment of engram-labelled (mCherry+) CA1 engram cell. Red arrowhead, mCherry+ subcellular compartment. Yellow arrowhead, microtubule. Purple arrowhead, mitochondrion. White arrowhead, knife damage from cryo-sectioning. Scale bars 100 nm. **D.** CryoET of unlabelled (ChR2-EYFP-/mCherry-) extracellular vesicle in engram-labelled mouse brain tissue. Top, tomographic slice. Dark red, extracellular vesicle. Bottom, ribosomes in a non-synaptic mCherry+ engram-labelled compartment. Cyan arrowhead, ribosome. **E.** Membrane-bound organelles within an mCherry+ engram-labelled compartment. Dark blue arrowhead, membrane-bound organelle. Scale bar, 20 nm.

## Notes

### Competing Interest Statement

The authors have declared no competing interest.

## References

1. Semon, R. (1904). Die Mneme als erhaltendes Prinzip im Wechsel des organischen Geschehens (Wilhelm Engelmann).

2. Liu, X., Ramirez, S., Pang, P.T., Puryear, C.B., Govindarajan, A., Deisseroth, K., and Tonegawa, S. (2012). Optogenetic stimulation of a hippocampal engram activates fear memory recall. Nature 484, 381–385. 10.1038/nature11028.

3. Liu, X., Ramirez, S., and Tonegawa, S. (2014). Inception of a false memory by optogenetic manipulation of a hippocampal memory engram. Philos. Trans. R. Soc. B: Biol. Sci. 369, 20130142. 10.1098/rstb.2013.0142.

4. Josselyn, S.A., Köhler, S., and Frankland, P.W. (2015). Finding the engram. Nat. Rev. Neurosci. 16, 521–534. 10.1038/nrn4000.

5. Josselyn, S.A., and Tonegawa, S. (2020). Memory engrams: Recalling the past and imagining the future. Science 367. 10.1126/science.aaw4325.

6. Ortega-de-San-Luis, C., and Ryan, T.J. (2022). Understanding the physical basis of memory: Molecular mechanisms of the engram. J. Biol. Chem. 298, 101866. 10.1016/j.jbc.2022.101866.

7. Hebb, D.O. (1949). The Organization of Behaviour: A Neuropsychologoical Theory.

8. Kandel, E.R., and Tauc, L. (1965). Heterosynaptic facilitation in neurones of the abdominal ganglion of Aplysia depilans. J. Physiol. 181, 1–27. 10.1113/jphysiol.1965.sp007742.

9. Bliss, T.V.P., and Lømo, T. (1973). Long-lasting potentiation of synaptic transmission in the dentate area of the anaesthetized rabbit following stimulation of the perforant path. J. Physiol. 232, 331–356. 10.1113/jphysiol.1973.sp010273.

10. Frey, U., and Morris, R.G.M. (1997). Synaptic tagging and long-term potentiation. Nature 385, 533–536. 10.1038/385533a0.

11. Turrigiano, G.G., and Nelson, S.B. (2004). Homeostatic plasticity in the developing nervous system. Nat. Rev. Neurosci. 5, 97–107. 10.1038/nrn1327.

12. Greer, P.L., and Greenberg, M.E. (2008). From Synapse to Nucleus: Calcium-Dependent Gene Transcription in the Control of Synapse Development and Function. Neuron 59, 846–860. 10.1016/j.neuron.2008.09.002.

13. Mansilla, A., Jordán-Álvarez, S., Santana, E., Jarabo, P., Casas-Tintó, S., and Ferrús, A. (2018). Molecular mechanisms that change synapse number. J. Neurogenet. 32, 155–170. 10.1080/01677063.2018.1506781.

14. Nabavi, S., Fox, R., Proulx, C.D., Lin, J.Y., Tsien, R.Y., and Malinow, R. (2014). Engineering a memory with LTD and LTP. Nature 511, 348–352. 10.1038/nature13294.

15. Chklovskii, D.B., Mel, B.W., and Svoboda, K. (2004). Cortical rewiring and information storage. Nature 431, 782–788. 10.1038/nature03012.

16. Tonegawa, S., Liu, X., Ramirez, S., and Redondo, R. (2015). Memory Engram Cells Have Come of Age. Neuron 87, 918–931. 10.1016/j.neuron.2015.08.002.

17. Ryan, T.J., Roy, D.S., Pignatelli, M., Arons, A., and Tonegawa, S. (2015). Engram cells retain memory under retrograde amnesia. Science 348, 1007–1013. 10.1126/science.aaa5542.

18. Kim, W.B., and Cho, J.-H. (2017). Encoding of Discriminative Fear Memory by Input-Specific LTP in the Amygdala. Neuron 95, 1129–1146.e5. 10.1016/j.neuron.2017.08.004.

19. Abdou, K., Shehata, M., Choko, K., Nishizono, H., Matsuo, M., Muramatsu, S., and Inokuchi, K. (2018). Synapse-specific representation of the identity of overlapping memory engrams. Science 360, 1227–1231. 10.1126/science.aat3810.

20. Lee, C., Lee, B.H., Jung, H., Lee, C., Sung, Y., Kim, H., Kim, J., Shim, J.Y., Kim, J., Choi, D.I., et al. (2023). Hippocampal engram networks for fear memory recruit new synapses and modify pre-existing synapses in vivo. Curr. Biol. 33, 507–516.e3. 10.1016/j.cub.2022.12.038.

21. Choi, J.-H., Sim, S.-E., Kim, J., Choi, D.I., Oh, J., Ye, S., Lee, J., Kim, T., Ko, H.-G., Lim, C.- S., et al. (2018). Interregional synaptic maps among engram cells underlie memory formation. Science 360, 430–435. 10.1126/science.aas9204.

22. Redondo, R.L., Kim, J., Arons, A.L., Ramirez, S., Liu, X., and Tonegawa, S. (2014). Bidirectional switch of the valence associated with a hippocampal contextual memory engram. Nature 513, 426–430. 10.1038/nature13725.

23. Vetere, G., Kenney, J.W., Tran, L.M., Xia, F., Steadman, P.E., Parkinson, J., Josselyn, S.A., and Frankland, P.W. (2017). Chemogenetic Interrogation of a Brain-wide Fear Memory Network in Mice. Neuron 94, 363–374.e4. 10.1016/j.neuron.2017.03.037.

24. Ryan, T.J., Luis, C.O. de S., Pezzoli, M., and Sen, S. (2021). Engram cell connectivity: an evolving substrate for information storage. Curr. Opin. Neurobiol. 67, 215–227. 10.1016/j.conb.2021.01.006.

25. Ortega-de-San-Luis, C., Pezzoli, M., Urrieta, E., and Ryan, T.J. (2023). Engram cell connectivity as a mechanism for information encoding and memory function. Curr. Biol. 10.1016/j.cub.2023.10.074.

26. Choi, J.-H., Sim, S.-E., Kim, J., Choi, D.I., Oh, J., Ye, S., Lee, J., Kim, T., Ko, H.-G., Lim, C.- S., et al. (2018). Interregional synaptic maps among engram cells underlie memory formation. Science 360, 430–435. 10.1126/science.aas9204.

27. Gray, E.G. (1959). Axo-somatic and axo-dendritic synapses of the cerebral cortex: an electron microscope study. J. Anat. 93, 420–433.

28. Bourne, J.N., and Harris, K.M. (2008). Balancing Structure and Function at Hippocampal Dendritic Spines. Annu. Rev. Neurosci. 31, 47–67. 10.1146/annurev.neuro.31.060407.125646.

29. Kulik, Y.D., Watson, D.J., Cao, G., Kuwajima, M., and Harris, K.M. (2019). Structural plasticity of dendritic secretory compartments during LTP-induced synaptogenesis. eLife 8, e46356. 10.7554/elife.46356.

30. Santuy, A., Tomás-Roca, L., Rodríguez, J.-R., González-Soriano, J., Zhu, F., Qiu, Z., Grant, S.G.N., DeFelipe, J., and Merchan-Perez, A. (2020). Estimation of the number of synapses in the hippocampus and brain-wide by volume electron microscopy and genetic labeling. Sci. Rep. 10, 14014. 10.1038/s41598-020-70859-5.

31. Fernández-Busnadiego, R., Zuber, B., Maurer, U.E., Cyrklaff, M., Baumeister, W., and Lučić, V. (2010). Quantitative analysis of the native presynaptic cytomatrix by cryoelectron tomography. J Cell Biol 188, 145–156. 10.1083/jcb.200908082.

32. Zuber, B., and Lučić, V. (2019). Molecular architecture of the presynaptic terminal. Curr Opin Struc Biol 54, 129–138. 10.1016/j.sbi.2019.01.008.

33. Radecke, J., Seeger, R., Kádková, A., Laugks, U., Khosrozadeh, A., Goldie, K.N., Lučić, V., Sørensen, J.B., and Zuber, B. (2023). Morphofunctional changes at the active zone during synaptic vesicle exocytosis. EMBO Rep. 24, EMBR202255719. 10.15252/embr.202255719.

34. Peukes, J., Lovatt, C., Leistner, C., Boulanger, J., Morado, D., Fuller, M., Kukulski, W., Zhu, F., Komiyama, N., Briggs, J., et al. (2024). The molecular infrastructure of glutamatergic synapses in the mammalian forebrain. 10.7554/elife.100335.

35. Tao, C.-L., Liu, Y.-T., Sun, R., Zhang, B., Qi, L., Shivakoti, S., Tian, C.-L., Zhang, P., Lau, P.-M., Zhou, Z.H., et al. (2018). Differentiation and Characterization of Excitatory and Inhibitory Synapses by Cryo-electron Tomography and Correlative Microscopy. J Neurosci 38, 1493– 1510. 10.1523/jneurosci.1548-17.2017.

36. Held, R.G., Liang, J., and Brunger, A.T. (2024). Nanoscale architecture of synaptic vesicles and scaffolding complexes revealed by cryo-electron tomography. Proc. Natl. Acad. Sci. 121, e2403136121. 10.1073/pnas.2403136121.

37. Held, R.G., Liang, J., Esquivies, L., Khan, Y.A., Wang, C., Azubel, M., and Brunger, A.T. (2024). In-Situ Structure and Topography of AMPA Receptor Scaffolding Complexes Visualized by CryoET. bioRxiv, 2024.10.19.619226. 10.1101/2024.10.19.619226.

38. Zuber, B., Nikonenko, I., Klauser, P., Muller, D., and Dubochet, J. (2005). The mammalian central nervous synaptic cleft contains a high density of periodically organized complexes. Proc. Natl. Acad. Sci. 102, 19192–19197. 10.1073/pnas.0509527102.

39. Matsui, A., Spangler, C., Elferich, J., Shiozaki, M., Jean, N., Zhao, X., Qin, M., Zhong, H., Yu, Z., and Gouaux, E. (2024). Cryo-electron tomographic investigation of native hippocampal glutamatergic synapses. eLife 13, RP98458. 10.7554/elife.98458.

40. Glynn, C., Smith, J.L.R., Case, M., Csöndör, R., Katsini, A., Sanita, M.E., Glen, T.S., Pennington, A., and Grange, M. (2024). Charting the molecular landscape of neuronal organisation within the hippocampus using cryo electron tomography. bioRxiv, 2024.10.14.617844. 10.1101/2024.10.14.617844.

41. Kukulski, W., Schorb, M., Welsch, S., Picco, A., Kaksonen, M., and Briggs, J.A.G. (2011). Correlated fluorescence and 3D electron microscopy with high sensitivity and spatial precision. J. Cell Biol. 192, 111–119. 10.1083/jcb.201009037.

42. Nakashiba, T., Young, J.Z., McHugh, T.J., Buhl, D.L., and Tonegawa, S. (2008). Transgenic Inhibition of Synaptic Transmission Reveals Role of CA3 Output in Hippocampal Learning. Science 319, 1260–1264. 10.1126/science.1151120.

43. Micheva, K.D., Busse, B., Weiler, N.C., O’Rourke, N., and Smith, S.J. (2010). Single-Synapse Analysis of a Diverse Synapse Population: Proteomic Imaging Methods and Markers. Neuron 68, 639–653. 10.1016/j.neuron.2010.09.024.

44. Castejón, O.J., Fuller, L., and Dailey, M.E. (2004). Localization of synapsin-I and PSD-95 in developing postnatal rat cerebellar cortex. Dev. Brain Res. 151, 25–32. 10.1016/j.devbrainres.2004.03.019.

45. Frank, R.A.W., Komiyama, N.H., Ryan, T.J., Zhu, F., Dell, T.J.O. rsquo, and Grant, S.G.N. (2016). NMDA receptors are selectively partitioned into complexes and supercomplexes during synapse maturation. Nature Communications 7, 11264.

46. Raisman, G., Cowan, W.M., and Powell, T.P.S. (1966). An experimental analysis of the efferent projection of the hippocampus. Brain 89, 83–108. 10.1093/brain/89.1.83.

47. Ishizuka, N., Weber, J., and Amaral, D.G. (1990). Organization of intrahippocampal projections originating from CA3 pyramidal cells in the rat. J. Comp. Neurol. 295, 580–623. 10.1002/cne.902950407.

48. Li, X. -G., Somogyi, P., Ylinen, A., and Buzsáki, G. (1994). The hippocampal CA3 network: An in vivo intracellular labeling study. J. Comp. Neurol. 339, 181–208. 10.1002/cne.903390204.

49. Shepherd, G.M.G., and Harris, K.M. (1998). Three-Dimensional Structure and Composition of CA3→CA1 Axons in Rat Hippocampal Slices: Implications for Presynaptic Connectivity and Compartmentalization. J. Neurosci. 18, 8300–8310. 10.1523/jneurosci.18-20-08300.1998.

50. Al-Amoudi, A., Norlen, L.P.O., and Dubochet, J. (2004). Cryo-electron microscopy of vitreous sections of native biological cells and tissues. J Struct Biol 148, 131–135. 10.1016/j.jsb.2004.03.010.

51. Wolf, S.G., Mutsafi, Y., Dadosh, T., Ilani, T., Lansky, Z., Horowitz, B., Rubin, S., Elbaum, M., and Fass, D. (2017). 3D visualization of mitochondrial solid-phase calcium stores in whole cells. eLife 6, e29929. 10.7554/elife.29929.

52. Delgado, T., Petralia, R.S., Freeman, D.W., Sedlacek, M., Wang, Y.-X., Brenowitz, S.D., Sheu, S.-H., Gu, J.W., Kapogiannis, D., Mattson, M.P., et al. (2019). Comparing 3D ultrastructure of presynaptic and postsynaptic mitochondria. Biol. Open 8, bio044834. 10.1242/bio.044834.

53. Knott, G.W., Quairiaux, C., Genoud, C., and Welker, E. (2002). Formation of Dendritic Spines with GABAergic Synapses Induced by Whisker Stimulation in Adult Mice. Neuron 34, 265–273. 10.1016/s0896-6273(02)00663-3.

54. Tamada, H., Blanc, J., Korogod, N., Petersen, C.C., and Knott, G.W. (2020). Ultrastructural comparison of dendritic spine morphology preserved with cryo and chemical fixation. eLife 9, e56384. 10.7554/elife.56384.

55. Yuste, R., and Bonhoeffer, T. (2004). Genesis of dendritic spines: insights from ultrastructural and imaging studies. Nat. Rev. Neurosci. 5, 24–34. 10.1038/nrn1300.

56. Chiu, C.Q., Lur, G., Morse, T.M., Carnevale, N.T., Ellis-Davies, G.C.R., and Higley, M.J. (2013). Compartmentalization of GABAergic Inhibition by Dendritic Spines. Science 340, 759– 762. 10.1126/science.1234274.

57. Konietzny, A., González-Gallego, J., Bär, J., Perez-Alvarez, A., Drakew, A., Demmers, J.A.A., Dekkers, D.H.W., Hammer, J.A., Frotscher, M., Oertner, T.G., et al. (2019). Myosin V regulates synaptopodin clustering and localization in the dendrites of hippocampal neurons. J. Cell Sci. 132, jcs230177. 10.1242/jcs.230177.

58. Bucher, M., Fanutza, T., and Mikhaylova, M. (2020). Cytoskeletal makeup of the synapse: Shaft versus spine. Cytoskeleton 77, 55–64. 10.1002/cm.21583.

59. Südhof, T.C. (2012). The Presynaptic Active Zone. Neuron 75, 11–25. 10.1016/j.neuron.2012.06.012.

60. Akert, K., and Sandri, C. (1968). An electron-microscopic study of zinc iodide-osmium impregnation of neurons. I. Staining of synaptic vesicles at cholinergic junctions. Brain Res. 7, 286–295. 10.1016/0006-8993(68)90104-2.

61. Borges-Merjane, C., Kim, O., and Jonas, P. (2020). Functional Electron Microscopy, “Flash and Freeze,” of Identified Cortical Synapses in Acute Brain Slices. Neuron 105, 992–1006.e6. 10.1016/j.neuron.2019.12.022.

62. Imig, C., López-Murcia, F.J., Maus, L., García-Plaza, I.H., Mortensen, L.S., Schwark, M., Schwarze, V., Angibaud, J., Nägerl, U.V., Taschenberger, H., et al. (2020). Ultrastructural Imaging of Activity-Dependent Synaptic Membrane-Trafficking Events in Cultured Brain Slices. Neuron 108, 843–860.e8. 10.1016/j.neuron.2020.09.004.

63. Papantoniou, C., Laugks, U., Betzin, J., Capitanio, C., Ferrero, J.J., Sánchez-Prieto, J., Schoch, S., Brose, N., Baumeister, W., Cooper, B.H., et al. (2023). Munc13- and SNAP25-dependent molecular bridges play a key role in synaptic vesicle priming. Sci. Adv. 9, eadf6222. 10.1126/sciadv.adf6222.

64. Okamoto, K.-I., Nagai, T., Miyawaki, A., and Hayashi, Y. (2004). Rapid and persistent modulation of actin dynamics regulates postsynaptic reorganization underlying bidirectional plasticity. Nat. Neurosci. 7, 1104–1112. 10.1038/nn1311.

65. Korobova, F., and Svitkina, T. (2010). Molecular Architecture of Synaptic Actin Cytoskeleton in Hippocampal Neurons Reveals a Mechanism of Dendritic Spine Morphogenesis. Mol. Biol. Cell 21, 165–176. 10.1091/mbc.e09-07-0596.

66. Basu, S., and Lamprecht, R. (2018). The Role of Actin Cytoskeleton in Dendritic Spines in the Maintenance of Long-Term Memory. Front. Mol. Neurosci. 11, 143. 10.3389/fnmol.2018.00143.

67. Goto, A., Bota, A., Miya, K., Wang, J., Tsukamoto, S., Jiang, X., Hirai, D., Murayama, M., Matsuda, T., McHugh, T.J., et al. (2021). Stepwise synaptic plasticity events drive the early phase of memory consolidation. Science 374, 857–863. 10.1126/science.abj9195.

68. Fukazawa, Y., Saitoh, Y., Ozawa, F., Ohta, Y., Mizuno, K., and Inokuchi, K. (2003). Hippocampal LTP Is Accompanied by Enhanced F-Actin Content within the Dendritic Spine that Is Essential for Late LTP Maintenance In Vivo. Neuron 38, 447–460. 10.1016/s0896-6273(03)00206-x.

69. Dimchev, G., Amiri, B., Fäßler, F., Falcke, M., and Schur, F.K. (2021). Computational toolbox for ultrastructural quantitative analysis of filament networks in cryo-ET data. J. Struct. Biol. 213, 107808. 10.1016/j.jsb.2021.107808.

70. Savtchenko, L.P., and Rusakov, D.A. (2007). The optimal height of the synaptic cleft. Proceedings Of The National Academy Of Sciences Of The United States Of America 104, 1823–1828.

71. Peters, A., Palay, P.S.L., and Webster, H.D.F. (1991). The fine structure of the nervous system: the neurons and supporting cells (Oxford University Press).

72. Briggs, J.A. (2013). Structural biology in situ—the potential of subtomogram averaging. Curr. Opin. Struct. Biol. 23, 261–267. 10.1016/j.sbi.2013.02.003.

73. Heumann, J.M., Hoenger, A., and Mastronarde, D.N. (2011). Clustering and variance maps for cryo-electron tomography using wedge-masked differences. J. Struct. Biol. 175, 288–299. 10.1016/j.jsb.2011.05.011.

74. Gilbert, M.A.G., Fatima, N., Jenkins, J., O’Sullivan, T.J., Schertel, A., Halfon, Y., Wilkinson, M., Morrema, T.H.J., Geibel, M., Read, R.J., et al. (2024). CryoET of β-amyloid and tau within postmortem Alzheimer’s disease brain. Nature 631, 913–919. 10.1038/s41586-024-07680-x.

75. Chang, S., and Camilli, P.D. (2001). Glutamate regulates actin-based motility in axonal filopodia. Nat. Neurosci. 4, 787–793. 10.1038/90489.

76. Menna, E., Disanza, A., Cagnoli, C., Schenk, U., Gelsomino, G., Frittoli, E., Hertzog, M., Offenhauser, N., Sawallisch, C., Kreienkamp, H.-J., et al. (2009). Eps8 Regulates Axonal Filopodia in Hippocampal Neurons in Response to Brain-Derived Neurotrophic Factor (BDNF). PLoS Biol. 7, e1000138. 10.1371/journal.pbio.1000138.

77. Ketschek, A., and Gallo, G. (2010). Nerve Growth Factor Induces Axonal Filopodia through Localized Microdomains of Phosphoinositide 3-Kinase Activity That Drive the Formation of Cytoskeletal Precursors to Filopodia. J. Neurosci. 30, 12185–12197. 10.1523/jneurosci.1740-10.2010.

78. Griswold, J.M., Bonilla-Quintana, M., Pepper, R., Lee, C.T., Raychaudhuri, S., Ma, S., Gan, Q., Syed, S., Zhu, C., Bell, M., et al. (2024). Membrane mechanics dictate axonal pearls-on-a-string morphology and function. Nat. Neurosci., 1–13. 10.1038/s41593-024-01813-1.

79. Leistner, C., Wilkinson, M., Burgess, A., Lovatt, M., Goodbody, S., Xu, Y., Deuchars, S., Radford, S.E., Ranson, N.A., and Frank, R.A.W. (2023). The in-tissue molecular architecture of β-amyloid pathology in the mammalian brain. Nat. Commun. 14, 2833. 10.1038/s41467-023-38495-5.

80. Rao-Ruiz, P., Yu, J., Kushner, S.A., and Josselyn, S.A. (2019). Neuronal competition: microcircuit mechanisms define the sparsity of the engram. Curr. Opin. Neurobiol. 54, 163–170. 10.1016/j.conb.2018.10.013.

81. Martinez-Sanchez, A., Laugks, U., Kochovski, Z., Papantoniou, C., Zinzula, L., Baumeister, W., and Lučić, V. (2021). Trans-synaptic assemblies link synaptic vesicles and neuroreceptors. Sci Adv 7, eabe6204. 10.1126/sciadv.abe6204.

82. Lovatt, M., Leistner, C., and Frank, R.A.W. (2022). Bridging length scales from molecules to the whole organism by cryoCLEM and cryoET. Faraday Discuss. 240, 114–126. 10.1039/d2fd00081d.

83. Uytiepo, M., Zhu, Y., Bushong, E., Polli, F., Chou, K., Zhao, E., Kim, C., Luu, D., Chang, L., Quach, T., et al. (2024). Synaptic architecture of a memory engram in the mouse hippocampus. bioRxiv, 2024.04.23.590812. 10.1101/2024.04.23.590812.

84. Zhu, F., Cizeron, M., Qiu, Z., Benavides-Piccione, R., Kopanitsa, M.V., Skene, N.G., Koniaris, B., DeFelipe, J., Fransén, E., Komiyama, N.H., et al. (2018). Architecture of the Mouse Brain Synaptome. Neuron 99, 781–799.e10. 10.1016/j.neuron.2018.07.007.

85. Bulovaite, E., Qiu, Z., Kratschke, M., Zgraj, A., Fricker, D.G., Tuck, E.J., Gokhale, R., Koniaris, B., Jami, S.A., Merino-Serrais, P., et al. (2022). A brain atlas of synapse protein lifetime across the mouse lifespan. Neuron 110, 4057–4073.e8. 10.1016/j.neuron.2022.09.009.

86. Wu, G.-H., Mitchell, P.G., Galaz-Montoya, J.G., Hecksel, C.W., Sontag, E.M., Gangadharan, V., Marshman, J., Mankus, D., Bisher, M.E., Lytton-Jean, A.K.R., et al. (2020). Multi-scale 3D Cryo-Correlative Microscopy for Vitrified Cells. Structure 28, 1231–1237.e3. 10.1016/j.str.2020.07.017.

87. Hebscher, M., Wing, E., Ryan, J., and Gilboa, A. (2019). Rapid Cortical Plasticity Supports Long-Term Memory Formation. Trends Cogn. Sci. 23, 989–1002. 10.1016/j.tics.2019.09.009.

88. Zhu, Y., Uytiepo, M., Bushong, E., Haberl, M., Beutter, E., Scheiwe, F., Zhang, W., Chang, L., Luu, D., Chui, B., et al. (2021). Nanoscale 3D EM reconstructions reveal intrinsic mechanisms of structural diversity of chemical synapses. Cell Rep. 35, 108953. 10.1016/j.celrep.2021.108953.

89. Roy, D.S., Park, Y.-G., Kim, M.E., Zhang, Y., Ogawa, S.K., DiNapoli, N., Gu, X., Cho, J.H., Choi, H., Kamentsky, L., et al. (2022). Brain-wide mapping reveals that engrams for a single memory are distributed across multiple brain regions. Nat. Commun. 13, 1799. 10.1038/s41467-022-29384-4.

90. Marco, A., Meharena, H.S., Dileep, V., Raju, R.M., Davila-Velderrain, J., Zhang, A.L., Adaikkan, C., Young, J.Z., Gao, F., Kellis, M., et al. (2020). Mapping the epigenomic and transcriptomic interplay during memory formation and recall in the hippocampal engram ensemble. Nat. Neurosci. 23, 1606–1617. 10.1038/s41593-020-00717-0.

91. Pignatelli, M., Ryan, T.J., Roy, D.S., Lovett, C., Smith, L.M., Muralidhar, S., and Tonegawa, S. (2019). Engram Cell Excitability State Determines the Efficacy of Memory Retrieval. Neuron 101, 274–284.e5. 10.1016/j.neuron.2018.11.029.

92. Park, S., Ko, S.Y., Frankland, P.W., and Josselyn, S.A. (2024). Comparing behaviours induced by natural memory retrieval and optogenetic reactivation of an engram ensemble in mice. Philos. Trans. B 379, 20230227. 10.1098/rstb.2023.0227.

93. Mocle, A.J., Ramsaran, A.I., Jacob, A.D., Rashid, A.J., Luchetti, A., Tran, L.M., Richards, B.A., Frankland, P.W., and Josselyn, S.A. (2024). Excitability mediates allocation of pre-configured ensembles to a hippocampal engram supporting contextual conditioned threat in mice. Neuron 112, 1487–1497.e6. 10.1016/j.neuron.2024.02.007.

94. Autore, L., O’Leary, J.D., Ortega-de-San-Luis, C., and Ryan, T.J. (2023). Adaptive expression of engrams by retroactive interference. Cell Rep. 42, 112999. 10.1016/j.celrep.2023.112999.

95. Ryan, T.J., and Frankland, P.W. (2022). Forgetting as a form of adaptive engram cell plasticity. Nat Rev Neurosci 23, 173–186. 10.1038/s41583-021-00548-3.

96. Denny, C.A., Lebois, E., and Ramirez, S. (2017). From Engrams to Pathologies of the Brain. Front. Neural Circuits 11, 23. 10.3389/fncir.2017.00023.

97. Power, S.D., Stewart, E., Zielke, L.G., Byrne, E.P., Douglas, A., San-Luis, C.O., Lynch, L., and Ryan, T.J. (2023). Immune activation state modulates infant engram expression across development. Sci. Adv. 9, eadg9921. 10.1126/sciadv.adg9921.

98. Chapman, D.P., Power, S.D., Vicini, S., Ryan, T.J., and Burns, M.P. (2024). Amnesia after Repeated Head Impact Is Caused by Impaired Synaptic Plasticity in the Memory Engram. J. Neurosci. 44, e1560232024. 10.1523/jneurosci.1560-23.2024.

99. Brosens, N., Lesuis, S.L., Rao-Ruiz, P., Oever, M.C. van den, and Krugers, H.J. (2024). Shaping Memories via Stress: A Synaptic Engram Perspective. Biol. Psychiatry 95, 721–731. 10.1016/j.biopsych.2023.11.008.

100. Li, J., Jiang, R.Y., Arendt, K.L., Hsu, Y.-T., Zhai, S.R., and Chen, L. (2020). Defective memory engram reactivation underlies impaired fear memory recall in Fragile X syndrome. eLife 9, e61882. 10.7554/elife.61882.

101. Li, M., Yang, X.-K., Yang, J., Li, T.-X., Cui, C., Peng, X., Lei, J., Ren, K., Ming, J., Zhang, P., et al. (2024). Ketamine ameliorates post-traumatic social avoidance by erasing the traumatic memory encoded in VTA-innervated BLA engram cells. Neuron 112, 3192–3210.e6. 10.1016/j.neuron.2024.06.026.

102. Whitaker, L.R., and Hope, B.T. (2018). Chasing the addicted engram: identifying functional alterations in Fos-expressing neuronal ensembles that mediate drug-related learned behavior. Learn. Mem. 25, 455–460. 10.1101/lm.046698.117.

103. Longueville, S., Nakamura, Y., Brami-Cherrier, K., Coura, R., Hervé, D., and Girault, J. (2021). Long-lasting tagging of neurons activated by seizures or cocaine administration in Egr1- CreERT2 transgenic mice. Eur. J. Neurosci. 53, 1450–1472. 10.1111/ejn.15060.

104. Lesuis, S.L., Park, S., Hoorn, A., Rashid, A.J., Mocle, A.J., Salter, E.W., Vislavski, S., Gray, M.T., Torelli, A.M., DeCristofaro, A., et al. (2024). Stress disrupts engram ensembles in lateral amygdala to generalize threat memory in mice. Cell. 10.1016/j.cell.2024.10.034.

105. Perusini, J.N., Cajigas, S.A., Cohensedgh, O., Lim, S.C., Pavlova, I.P., Donaldson, Z.R., and Denny, C.A. (2017). Optogenetic stimulation of dentate gyrus engrams restores memory in Alzheimer’s disease mice. Hippocampus 27, 1110–1122. 10.1002/hipo.22756.

106. Poll, S., Mittag, M., Musacchio, F., Justus, L.C., Giovannetti, E.A., Steffen, J., Wagner, J., Zohren, L., Schoch, S., Schmidt, B., et al. (2020). Memory trace interference impairs recall in a mouse model of Alzheimer’s disease. Nat. Neurosci. 23, 952–958. 10.1038/s41593-020-0652-4.

107. Ting, J.T., Lee, B.R., Chong, P., Soler-Llavina, G., Cobbs, C., Koch, C., Zeng, H., and Lein, E. (2018). Preparation of Acute Brain Slices Using an Optimized N-Methyl-D-glucamine Protective Recovery Method. J. Vis. Exp. : JoVE, 53825. 10.3791/53825.

108. Wakayama, S., Kiyonaka, S., Arai, I., Kakegawa, W., Matsuda, S., Ibata, K., Nemoto, Y.L., Kusumi, A., Yuzaki, M., and Hamachi, I. (2017). Chemical labelling for visualizing native AMPA receptors in live neurons. Nat. Commun. 8, 14850. 10.1038/ncomms14850.

109. Studer, D., Klein, A., Iacovache, I., Gnaegi, H., and Zuber, B. (2014). A new tool based on two micromanipulators facilitates the handling of ultrathin cryosection ribbons. J Struct Biol 185, 125–128. 10.1016/j.jsb.2013.11.005.

110. Schorb, M., and Briggs, J.A.G. (2014). Correlated cryo-fluorescence and cryo-electron microscopy with high spatial precision and improved sensitivity. Ultramicroscopy 143, 24–32. 10.1016/j.ultramic.2013.10.015.

111. Kremer, J.R., Mastronarde, D.N., and McIntosh, J.R. (1996). Computer Visualization of Three-Dimensional Image Data Using IMOD. J. Struct. Biol. 116, 71–76. 10.1006/jsbi.1996.0013.

112. Liu, Y.-T., Zhang, H., Wang, H., Tao, C.-L., Bi, G.-Q., and Zhou, Z.H. (2022). Isotropic reconstruction for electron tomography with deep learning. Nat. Commun. 13, 6482. 10.1038/s41467-022-33957-8.

113. Navarro, P.P., Stahlberg, H., and Castaño-Díez, D. (2018). Protocols for Subtomogram Averaging of Membrane Proteins in the Dynamo Software Package. Front. Mol. Biosci. 5, 82. 10.3389/fmolb.2018.00082.

114. Pettersen, E.F., Goddard, T.D., Huang, C.C., Meng, E.C., Couch, G.S., Croll, T.I., Morris, J.H., and Ferrin, T.E. (2021). UCSF ChimeraX: Structure visualization for researchers, educators, and developers. Protein Sci. 30, 70–82. 10.1002/pro.3943.

115. Caspar, D.L.D., and Kirschner, D.A. (1971). Myelin Membrane Structure at 10 Å Resolution. Nat. N. Biol. 231, 46–52. 10.1038/newbio231046a0.

116. Mattila, P.K., and Lappalainen, P. (2008). Filopodia: molecular architecture and cellular functions. Nat. Rev. Mol. Cell Biol. 9, 446–454. 10.1038/nrm2406.

117. Fujii, T., Iwane, A.H., Yanagida, T., and Namba, K. (2010). Direct visualization of secondary structures of F-actin by electron cryomicroscopy. Nature 467, 724–728. 10.1038/nature09372.

